# Evaluating Study Design Rigor in Preclinical Cardiovascular Research: A Replication Study

**DOI:** 10.1101/2023.06.27.546731

**Authors:** Isaiah C. Jimenez, Gabrielle C. Montenegro, Keyana Zahiri, Damini Patel, Adrienne Mueller

## Abstract

**Background:** Methodological rigor remains a priority in preclinical cardiovascular research to ensure experimental reproducibility and high-quality research. Limited reproducibility diminishes the translation of preclinical discoveries into medical practice. In addition, lack of reproducibility fosters uncertainty in the public’s acceptance of reported research results.

**Methods:** We evaluated the reporting of methodological practices in preclinical cardiovascular research studies published in leading scientific journals by screening articles for the inclusion of the following study design elements (SDEs): considering sex as a biological variable, randomization, blinding, and sample size power estimation. We screened for these SDEs across articles regarding preclinical cardiovascular research studies published between 2011 and 2021. We replicated and extended a study published in 2017 by Ramirez et al. We hypothesized a higher SDE inclusion across preclinical studies over time, that preclinical studies that include human and animal substudies within the same study will exhibit greater SDE inclusion than animal-only preclinical studies, and that a difference exists in SDE usage between large and small animal models.

**Results:** SDE inclusion was low; with 15.2% of animal-only studies including both sexes as a biological variable, 30.4% including randomization, 32.1% including blinding, and 8.2% including sample size estimation. The incorporation of SDEs did not significantly increase over the ten-year timeframe in the screened articles. Randomization and sample size estimation differed significantly between animal and human substudies (corrected p=1.85e-05 and corrected p=3.81e-07, respectively.)

**Conclusions:** Evidence of methodological rigor varies depending on the study type and model organisms used. From 2011-2021, SDE reporting within preclinical studies has not increased, suggesting more work is needed to foster the inclusion of rigorous study design elements in cardiovascular research.

## INTRODUCTION

Preclinical studies using animal models play an important role in developing new treatments and evaluating the safety and efficacy of novel therapies. Preclinical cardiovascular research has greatly contributed to our understanding of heart disease (Houser et al., 2012; Bacmeister et al, 2019), yet we often find failed translations from “bench-to-bedside” (Justice & Dhillon, 2016; Seok et al., 2012). Methodological rigor in these studies remains a major priority to establish a level of consistent reproducibility across preclinical research.

One means to enhance reproducibility of preclinical research is to increase the frequency of use of study design elements (SDEs) such as inclusion of sex as a biological variable, randomization of samples or subjects, blinding, and sample size estimation. Including both biological sexes in experimental design removes sex as a potential confounding variable in establishing causal relationships between variables of interest. Implementation of randomization in experimental design is an additional means to control for *all* potential confounders between variables of interest. Moreover, randomization helps reduce bias. The use of blinding in experimental design limits potential bias in the assessment of experimental outcomes from study participants and researchers themselves. Finally, sample size estimation limits impractical significance in experimental results and false-negative results. Sample size estimation also addresses the ethical concern of over-sampling of animal model cohorts by establishing a minimum sample size needed. Ultimately, each of these four SDEs influences the reproducibility of preclinical experimental outcomes.

This study is a replication and extension of a study performed by Ramirez et al. (2017), which investigated the prevalence of these four SDEs in preclinical studies published in five leading cardiovascular journals of the American Heart Association (*Circulation*; *Circulation Research*; *Hypertension*; *Stroke*; and *Arteriosclerosis, Thrombosis, and Vascular Biology* (*ATVB*)) between July 2006 and June 2016. Their study found a low prevalence of SDEs across screened studies, reflecting low methodological rigor in preclinical cardiovascular research in that decade (Ramirez et al., 2017). This study investigates the inclusion of these four SDEs in randomly selected preclinical cardiovascular studies published between 2011 and 2021 in nine different leading biomedical and scientific journals outside of American Heart Association publications: *Science, Nature, European Heart Journal, Journal of the American College of Cardiology, New England Journal of Medicine, Cell, Lancet, Journal of the American Medical Association*, and *Proceedings of the National Academy of Sciences of the United States of America*. This study also examines the use of rigorous SDEs by comparing animal-only studies with studies that have both animal and human substudies (human/animal studies) over ten years. The decade 2011-2021 was selected for analysis to provide a more recent evaluation of methodological rigor in preclinical cardiovascular research, following the work of Ramirez et al., 2017.

By identifying trends in the sex of study subjects used, randomization, blinding, and sample size estimations, we assessed the methodological rigor of scientific practices carried out in preclinical cardiovascular research.

## METHODS

We reviewed preclinical cardiovascular articles published between 2011 and 2021 in leading biomedical and scientific journals. Studies were included from nine leading journals: *Science, Nature, European Heart Journal, Journal of the American College of Cardiology, New England Journal of Medicine, Cell, Lancet, Journal of the American Medical Association*, and *Proceedings of the National Academy of Sciences of the United States of America*. These journals were selected to complement those used in a previous study: *Circulation*; *Circulation Research*; *Hypertension*; *Stroke*; and *Arteriosclerosis, Thrombosis, and Vascular Biology* (*ATVB*) (Ramirez et al., 2017). Using a search string in a Pubmed query, we identified primary research articles (excluding editorials or comments) describing cardiovascular experiments using animal models. The complete search string used was:

((cardi*[Title]) OR (heart[Title]) OR (arteri*[Title]) OR (hypertensi*[Title]) OR (atherosclero*[Title]) OR (arrhythm*[Title])) AND ((pig[Title/Abstract]) OR (rat[Title/Abstract]) OR (mouse[Title/Abstract]) OR (guinea pig[Title/Abstract]) OR (gerbil[Title/Abstract]) OR (hamster[Title/Abstract]) OR (monkey[Title/Abstract]) OR (rabbit[Title/Abstract]) OR (dog[Title/Abstract])) NOT ((review[Publication Type]) OR (systematic review[Publication Type]) OR (editorial[Publication Type]) OR (comment[Publication Type])) AND ((“2011/01/01”[Date - Publication] : “2021/12/31”[Date - Publication])) AND ((“Lancet (London, England)”[Journal]) OR (“Nature”[Journal]) OR (“Science (New York, N.Y.)”[Journal]) OR (“JAMA”[Journal]) OR (“The New England journal of medicine”[Journal]) OR (“Proceedings of the National Academy of Sciences of the United States of America”[Journal]) OR (“Cell”[Journal]) OR (“European heart journal”[Journal]) OR (“Journal of the American College of Cardiology”[Journal])).

309 articles were returned by the PubMed Search query. No stopping rule was utilized, as the sample size was predetermined before data collection. Studies were included if they were published manuscripts and used animal subjects. Articles were excluded from data analysis if they did not include animal-model experiments, if they were not related to a cardiovascular research topic, or were published as an abstract, editorial, or any form other than a published full manuscript. Thus, although the search string yielded 309 studies for screening, 11 studies were excluded for not meeting inclusion criteria. A total of 298 studies were ultimately included in our data analyses.

Articles were screened based on four study design elements (SDEs): inclusion of both biological sexes in study subjects, randomization, blinding, and sample size estimation. The screening database included animal-only studies, as well as animal studies that included human substudies (human/animal studies). For human/animal studies, the same SDEs were used to evaluate methodological rigor across human subject populations. We evaluated the four SDEs separately for studies that only performed animal experiments and studies that performed human/animal experiments, if applicable. Screening definitions were predefined (Table I). Articles were also screened for what cardiovascular research topic they were investigating, as well as which animal species were used in the study (Table II). We initially used cardiovascular research topics that were defined by Ramirez et al. (2017) and also expanded the topic list to include 2 additional topics that occurred more frequently in our dataset: congenital heart disease and heart development/repair/regeneration.

**Table I:**
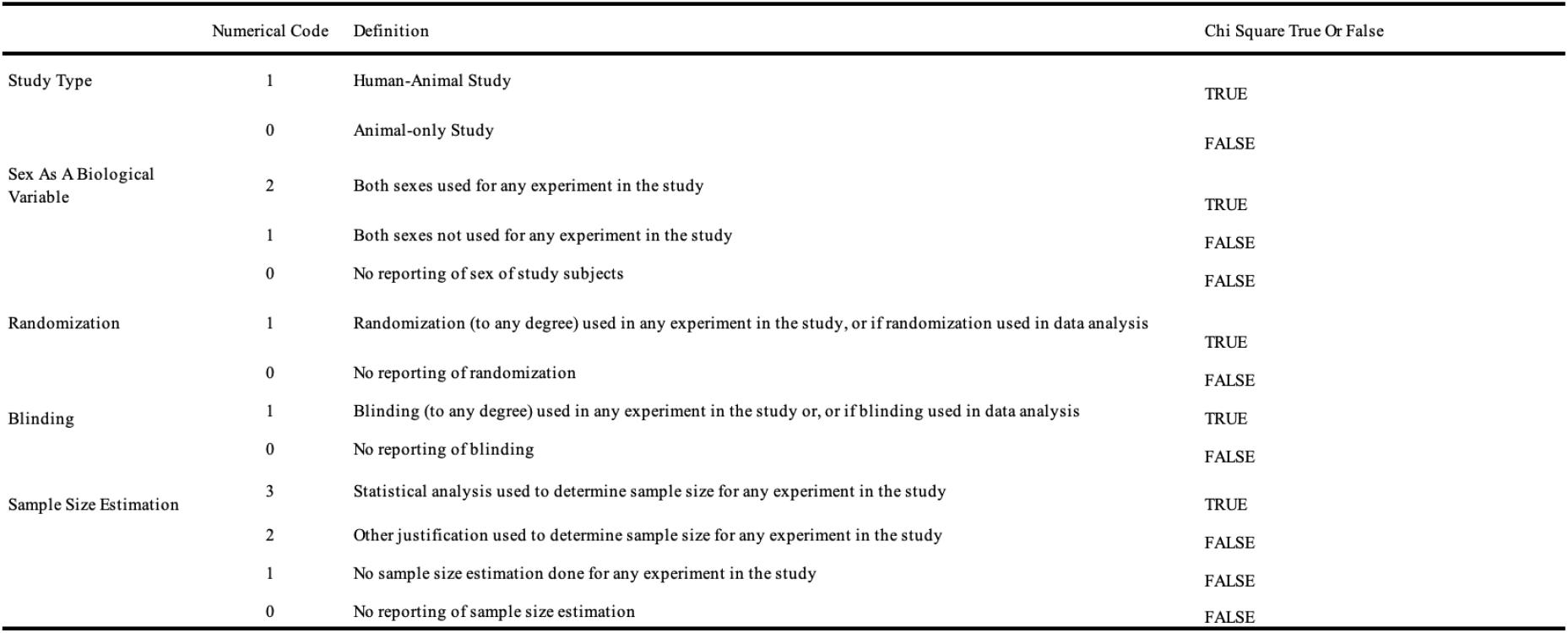
This table depicts the predetermined screening numerical codes and descriptions used for study analysis of methodological rigor across all studies.

**Table II:**
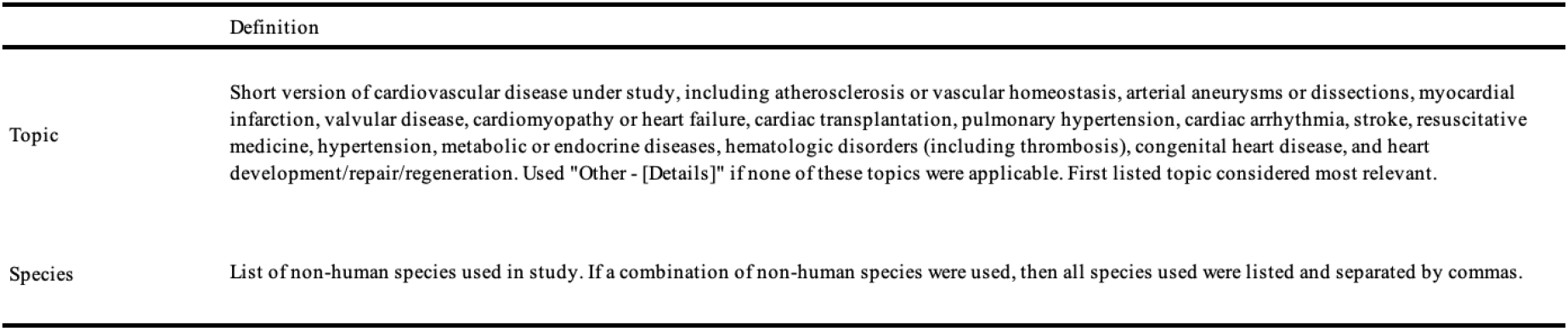
This table lists the information collected during screening across all studies.

Articles were distributed equally among members of the research team for screening. Articles were initially screened in the order they were returned from the PubMed search. Investigators did not select which articles from the database of 298 articles to screen based on any specific criteria. To ensure accuracy and consistency in screening, each article was independently screened by two investigators. Screeners were randomly assigned for the second screening of an article. Discrepancies in screening were resolved by consensus.

### Statistical Methods

The collected data was analyzed to evaluate the prevalence of the use of SDEs in preclinical cardiovascular research between 2011 and 2021. Categorical variables are reported as a number (%). RStudio and Microsoft Excel software were used to perform statistical significance comparisons. A median-based linear model analysis was performed to analyze changes in SDE inclusion over time, and nonparametric Kruskal-Wallis tests were performed to analyze differences in SDE reporting across journals. Chi-squared tests were performed for other analyses to assess differences in SDE reporting across experimental models, between animals vs humans within the same study, and across different animal models. A threshold of p < 0.05 was considered statistically significant, however, we used the Holm-Bonferroni correction to correct for multiple comparisons.

### Pre-registration and Data Availability

This study was pre-registered in the Open Science Framework (OSF) registry. The preregistration can be accessed via the following link: https://doi.org/10.17605/OSF.IO/F4NH9 (Patel et al., 2022). We adhered to the methodology detailed in the pre registration for this analysis.

The data and analytical methods used for this study are available at https://osf.io/52Q6W/ (Zahiri et al., 2022.)

## RESULTS

A total of 298 preclinical research studies published between 2011 and 2021 were included in our analyses. 61.7% (N=184) of these studies were animal only and 38.3% (N=114) were human/animal studies (Figure 1A). Approximately the same number of cardiovascular preclinical research studies were published for each of the ten years in our sample (Figure 1B). The majority of the studies in our analysis were published in Proceedings of the National Academy of Sciences of the United States of America, 45.3% (N=135) (Figure 1C). In addition, a wide range of species were used in the preclinical studies we analyzed (Figure 1D). Mice were the most commonly studied species used in 77.2% (N=230) of studies. This was followed by rats in 20.1% (N=60) of studies. Additionally, a wide range of topics relating to cardiovascular disease were investigated in the studies we assessed (Figure 1E). We categorized the topics as follows: cardiomyopathy or heart failure (28.2%), atherosclerosis or vascular homeostasis (16.1%), myocardial infarction (14.8%), cardiac arrhythmia (9.1%), heart development/repair/regeneration (8.0%), hypertension (4.4%), metabolic or endocrine disease (4.0%), congenital heart disease (2.7%), valvular disease (2.0%), cardiac transplantation (1.7%), hematological disorder (0.3%), and other (8.7%), based on topics identified in the original Ramirez et al. (2017) study.

**Figure 1:**
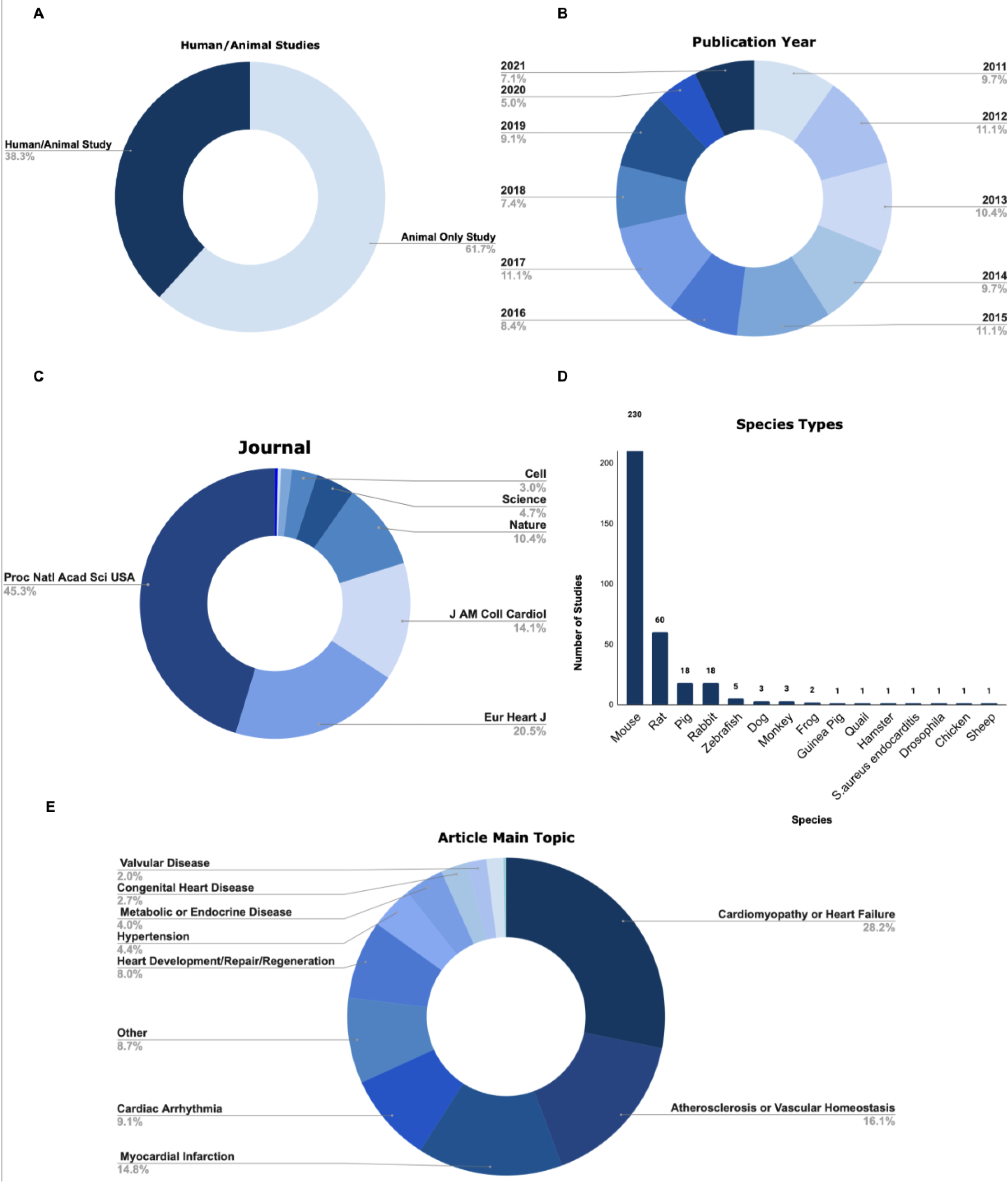
There is a diverse range of preclinical cardiovascular research studies published over the last decade. **(A)** This figure depicts the proportion of studies included in our analysis were animal only or human/animal studies. **(B)** This figure depicts the proportions of studies included in our analysis from various publication years between 2011-2021. (**C)** This figure depicts the proportions of studies included in our analysis from each of nine high impact scientific journals. (**D)** This figure depicts the range of species used in the sample populations of studies included in our analysis. Mice and rats were the most commonly studied animal models across all studies. (**E)** This figure depicts the main range of topics the articles used in this study primarily investigated.

### Overall SDE Inclusion

Table III shows the proportion of studies that included each of the four SDEs, stratified by animal-only studies, as well as animal and human substudies in human/animal studies. In animal-only studies, both sexes were used in 15.2% (N=28) studies, single-sex was used in 48.4% (N=89) studies, and there was no reporting on the sex of study subjects used in 36.4% (N=67) studies. In animal substudies in human/animal studies, both sexes were used in 17.5% (N=20) studies, single-sex was used in 53.5% (N=61) studies, and there was no reporting on the sex of study subjects used in 29.0% (N=33) studies. In human substudies in human/animal studies, both sexes were used in 36% (N=41) of studies, single-sex was used in 8.8% (N=10) of studies, and there was no reporting on the sex of study subjects used in 55.2% (N=63) studies.

**Table III:** These tables depict the descriptive statistics for SDEs analyzed for animal only studies vs animal or human substudies in human/animal studies.

| Both Sex Inclusion |  |  |  |  |
| --- | --- | --- | --- | --- |
|  | Both Sexes (N, %) | Single Sex (N, %) | Not Reported (N, %) |  |
| Animal Only | 28 (15.2) | 89 (48.4) | 67 (36.4) |  |
| Human/Animal - Animal Substudy | 20 (17.5) | 61 (53.5) | 33 (29) |  |
| Human/Animal - Human Substudy | 41 (36) | 10 (8.8) | 63 (55.2) |  |
| Randomization Inclusion |  |  |  |  |
|  | Randomization Used (N, %) | Randomization Not Used (N, %) |  |  |
| Animal Only | 56 (30.4) | 128 (69.6) |  |  |
| Human/Animal - Animal Substudy | 41 (36) | 73 (64) |  |  |
| Human/Animal - Human Substudy | 7 (6) | 107 (94) |  |  |
| Blinding Inclusion |  |  |  |  |
|  | Blinding Used (N, %) | Blinding Not Used (N, %) |  |  |
| Animal Only | 59 (32.1) | 125 (67.9) |  |  |
| Human/Animal - Animal Substudy | 28 (24.5) | 86 (75.5) |  |  |
| Human/Animal - Human Substudy | 13 (11.4) | 101 (88.6) |  |  |
| Sample Size Estimation Inclusion |  |  |  |  |
|  | Used Size Estimation (N, %) | Justified Not Using Size Estimation (N, %) | Size Estimation Not Used (N, %) | Not Reported (N, %) |
| Animal Only | 15 (8.2) | 9 (4.9) | 5 (2.7) | 155 (84.2) |
| Human/Animal - Animal Substudy | 8 (7) | 3 (2.6) | 6 (5.3) | 97 (85.1) |
| Human/Animal - Human Substudy | 2 (1.8) | 14 (12.3) | 3 (2.6) | 95 (83.3) |

In terms of randomization, 30.4% (N=56) of animal-only studies, 36% (N=41) of animal substudies in human/animal studies, and 6% (N=7) of human substudies in human/animal studies used randomization to any degree in their experiments. In terms of blinding, 32.1% (N=59) of animal-only studies, 24.5% (N=28) of animal substudies in human/animal studies, and 11.4% (N=13) of human substudies in human/animal studies used blinding to any degree in their experiments.

In terms of sample size estimations for animal-only studies, 8.2% (N=15) used statistical analysis to determine the sample size for experiments, 4.9% (N=9) provided other justification for the sample size they selected, 2.7% (N=5) indicated that no sample size estimation was done, and 84.2% (N=155) did not report sample size justification. Examples of ‘other justification for sample size selection’ include estimating sample size based on pilot studies or determining sample size based on previous standards in the field. In terms of sample size estimations for animal substudies in human/animal studies, 7% (N=8) used statistical analysis to determine the sample size for experiments, 2.6% (N=3) provided other justification for the sample size they selected, 5.3% (N=6) indicated that no sample size estimation was done, and 85.1% (N=97) did not report any sample size justification. Lastly, in terms of sample size estimations for human substudies in human/animal studies, 1.8% (N=2) used statistical analysis to determine the sample size for experiments, 12.3% (N=14) provided other justification for the sample size they selected, 2.6% (N=3) indicated that no sample size estimation was done, and 83.3% (N=95) did not report any sample size justification.

### Changes in SDEs Over Time

A median-based linear analysis of SDE inclusion between 2011 and 2021 revealed no statistically significant difference in the proportion of studies including animals of both biological sexes between 2011 and 2021 (corrected p=1.476) (Figure 2). Of studies screened from 2011, 10% (N=3 of 29) included animals of both biological sexes. Of studies screened from 2017, 27% (N=9 of 33) included animals of both biological sexes. This proportion ultimately decreases to 10% (N=2 of 21) of studies in 2021 (Figure 2). Although there appears to be an overall general increase in the proportion of studies including animals of both biological sexes across the decade of interest, no statistically significant difference was found (corrected p>0.05).

**Figure 2:**
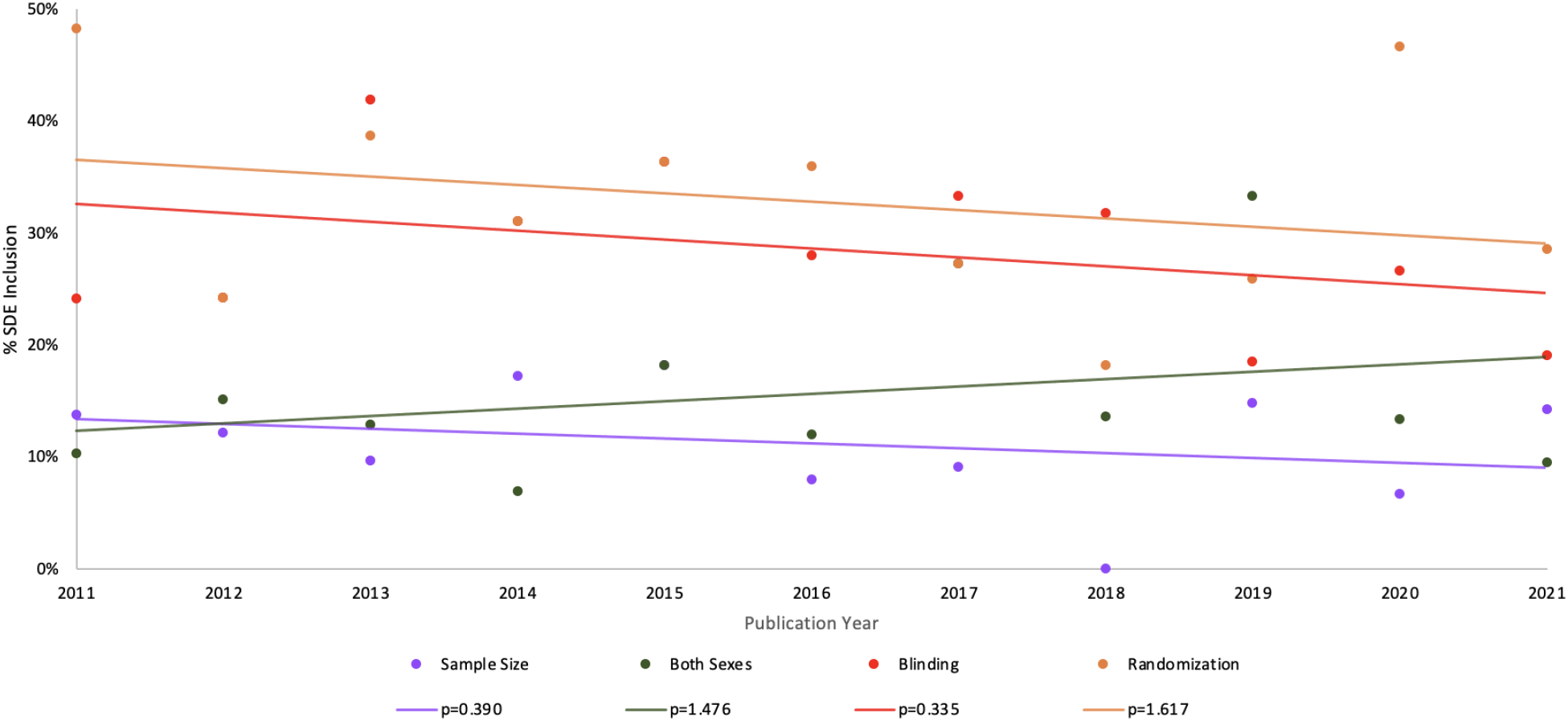
Percentage of studies with SDE inclusion in preclinical cardiovascular research between 2011-2021 differed across the four SDEs screened. Percentages are reported for animal models used in all 298 eligible studies, regardless of whether the experimental design included human substudies. The percentage of studies including both biological sexes, randomization, blinding, and sample size estimation showed no statistically significant difference between 2011-2021 (median-based linear analysis).

Similarly, a median-based linear analysis of randomization implementation revealed no statistically significant difference in the proportion of studies implementing randomization between 2011 and 2021 (corrected p=1.617). Approximately half of preclinical cardiovascular studies implemented randomization in 2011, as 48% (N=14 of 29) of 2011 studies mentioned randomization in their methods. This proportion declined to 29% (N=6 of 21) of studies in 2021 (Figure 2). Although there appears to be an overall general decrease in the proportion of studies including randomization across the decade of interest, no statistically significant decrease was found (corrected p>0.05)

Interestingly, a median-based linear analysis of blinding implementation also showed no statistically significant difference in the proportion of studies including blinding as part of the study design between 2011 and 2021 (corrected p=0.335) (Figure 2). Of studies screened from 2011, 24% (N=7 of 29) included blinding. This proportion decreased to 19% (N=4 of 21) in 2021 (Figure 2). Likewise, comparable results were found for sample size justification (corrected p=0.390) (Figure 2). Of studies screened from 2011, 14% (N=4 of 29) justified their sample size using size estimation (justification or statistical estimation). This proportion dropped to 0% (N=0 of 22) studies in 2018 but ultimately increased back up to 14% (N=3 of 21) in 2021 (Figure 2). Again, although there appears to be an overall general decrease in the proportion of studies including blinding and sample size justification as part of the study design, a median-based linear model ultimately revealed no statistically significant decrease across the decade of interest (corrected p>0.05).

### Differences in SDE Reporting Across Journals

Of all 298 journal articles screened in this study, the four journals with the highest numbers of cardiovascular preclinical studies were: *Proceedings of the National Academy of Sciences of the United States of America (Proc Natl Acad Sci)* 45.3% (N=135), *European Heart Journal (Eur Heart J)* 20.5% (N=61), *Journal of the American College of Cardiology (J Am Coll Cardiol)* 14.1% (N=42), and *Nature* 10.4% (N=31). A general difference in the reporting of the four SDEs was observed across all four journals, but differences were not determined to be statistically significant by Kruskal-Wallis tests: both sexes: H(10)=6.625, corrected p=2.281; randomization: H(10)=6.597, corrected p=1.526; blinding: H(10)=3.498, corrected p=0.967, sample size estimation: H(8)=8.344, corrected p=2.804) (Figure 3).

**Figure 3:**
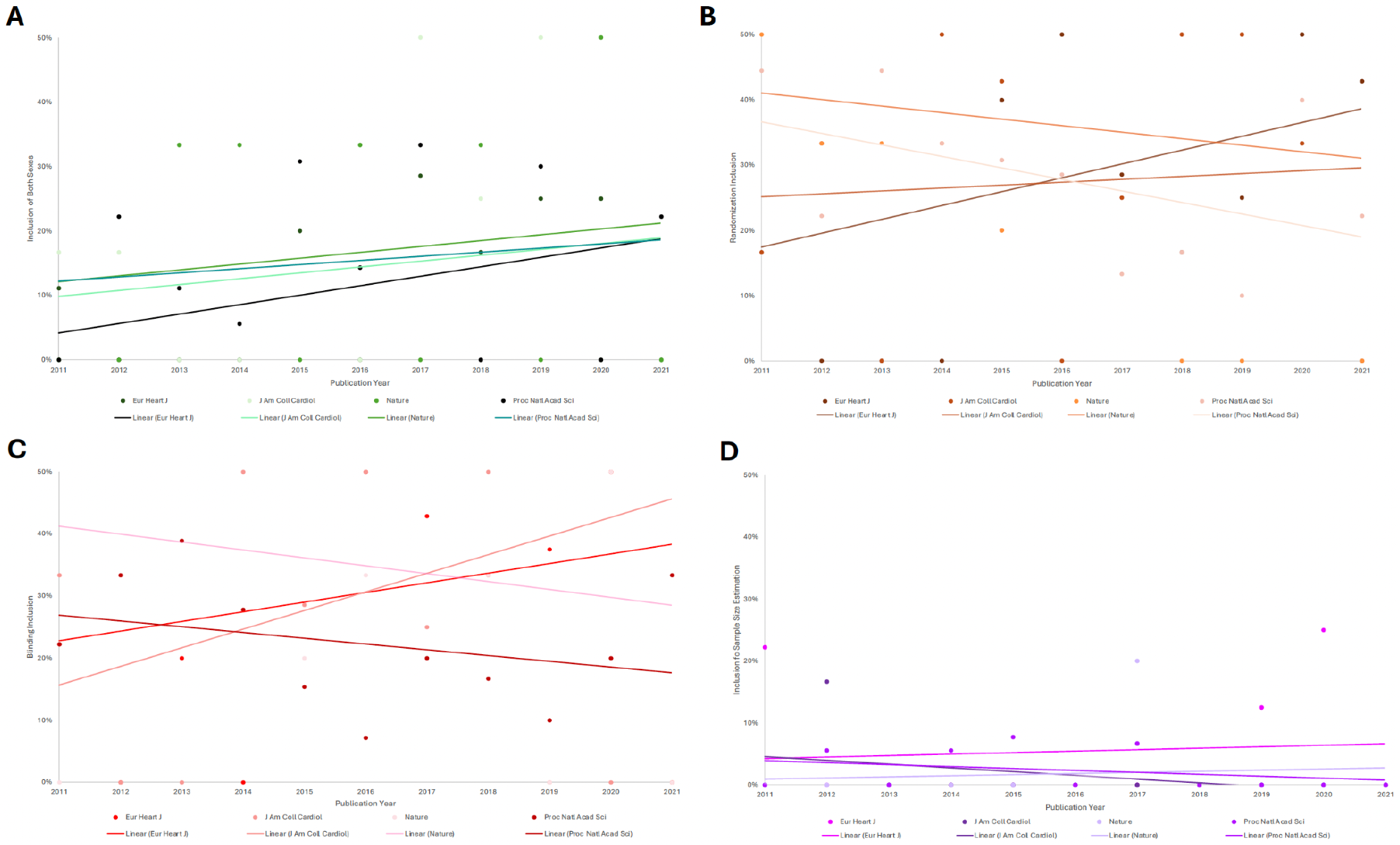
The percentage of SDE inclusion in preclinical cardiovascular research between 2011-2021 generally varied across the four most prevalent journals. **(A) Both Sexes Analysis:** The percentage of studies including animals of both biological sexes generally increased across all four most journals, but no statistically significant differences were found (corrected p=2.281) (Kruskal-Wallis test analysis). (**B) Randomization Analysis:** The percentage of studies using randomization generally increased in studies from *Eur Heart J* and *J Am Coll Cardiol*, but decreased across studies from *Proc Natl Acad Sci* and *Nature* (Kruskal-Wallis Test). However, no statistically significant differences were found (corrected p=1.526) **(C) Blinding Analysis:** Similar results for randomization inclusion apply to blinding inclusion. Likewise, no statistically significant differences were found (corrected p=0.967) (Kruskal-Wallis test analysis). (**D) Sample Size Analysis:** The percentage of studies implementing statistical sample size estimations increased in studies screened from *Eur Heart J* and *Nature*, and decreased across those screened from *Proc Natl Acad Sci* and *J Am Coll Cardiol* (Kruskal-Wallis Test analysis). Still, no statistically significant differences were found (corrected p=2.804).

Articles screened across the four most prevalent journals in this study varied in frequency across the ten years (2011-2021). Among studies screened from *Eur Heart J*, (N=1 of 11) 11% included animals of both biological sexes in 2011 while (N=2 of 8) 25% included both biological sexes in 2019. Across studies screened from the *J Am Coll Cardiol*, (N=1 of 6) 17% included animals from both biological sexes in 2011 while 50% (N=2 of 4) included both biological sexes in 2019 (Figure 3). A difference of +14% in the *Eur Heart J* and +33% in the *J Am Coll Cardiol* between the journals was not found to be statistically significant (corrected p=2.804).

In studies screened from *Eur Heart J*, 56% (N=5 of 9) reported using randomization in 2011 and 43% (N=3 of 7) reported using randomization in 2021. Meanwhile, in studies screened from the *Proc Natl Acad Sci*, 44% (N=4 of 9) reported using randomization in 2011 and 22% (N=2 of 9) reported using randomization in 2021 (Figure 3). Similarly, although a difference of -13% over time was observed in *Eur Heart J* versus a difference of -22% in the *Proc Natl Acad Sci*, no statistically significant difference between these decreasing rates was found across the ten years (corrected p=1.526).

With regards to reporting of blinding, in studies screened from the *Proc Natl Acad Sci*, 22% (N=2 of 9) reported implementing blinding in 2011 and 33% (N=3 of 9) reported implementing blinding in 2021. In studies screened from the *J Am Coll Cardiol*, 33% (N=2 of 6) used blinding in 2011 while 100% (N=1 of 1) used blinding in 2021 (Figure 3). A change of +11% overtime was observed across studies screened from the *Proc Natl Acad Sci* versus a change of +77% across studies screened from the *J Am Coll Cardiol*, but no statistically significant difference was found in comparing these differences as well (corrected p=0.967).

Finally, in assessing differences in reporting statistical estimations for study sample size, a difference of +3% (N=2 of 9 to N=1 of 4) was found across studies screened from the *Eur Heart J* between 2011 and 2020 (Figure 3). A difference of -10% (N=3 of 18 to N=1 of 15) was observed between studies screened from *Proc Natl Acad Sci* from 2012-2017 (Figure 3). No studies before 2012 nor beyond 2017 from the *Proc Natl Acad Sci* reported using sample size estimations. Observed differences were not determined to be statistically significant (corrected p=2.804).

### Differences in SDE Reporting Across Experimental Models

There were slight differences in SDE reporting for animal substudies in human/animal studies vs animal-only studies (Figure 4). In human/animal studies, 18% (N=20) used subjects of both sexes, 53% (N=61) used only one sex of study subjects, and 29% (N= 33) did not report the sex of study subjects used, as opposed to 16% (N=28), 47% (N=89), and 37% (N=67), respectively, for animal only studies. Randomization was only used in 36% (N=41) of human/animal studies and 30% (N=56) of animal-only studies. Blinding was only used in 25% (N=28) of human/animal studies and 32% (N=59) of animal-only studies. In terms of sample size estimations for human/animal studies, 7% (N=8) performed sample size estimations, 3% (N=3) justified not using sample size estimations, 5% (N=6) did not perform sample size estimation, and 85% (N=97) did not report any information on sample size estimation. For animal-only studies, these values were 8% (N=15), 5% (N=9), 3% (N=5), and 84% (N=155), respectively.

**Figure 4:**
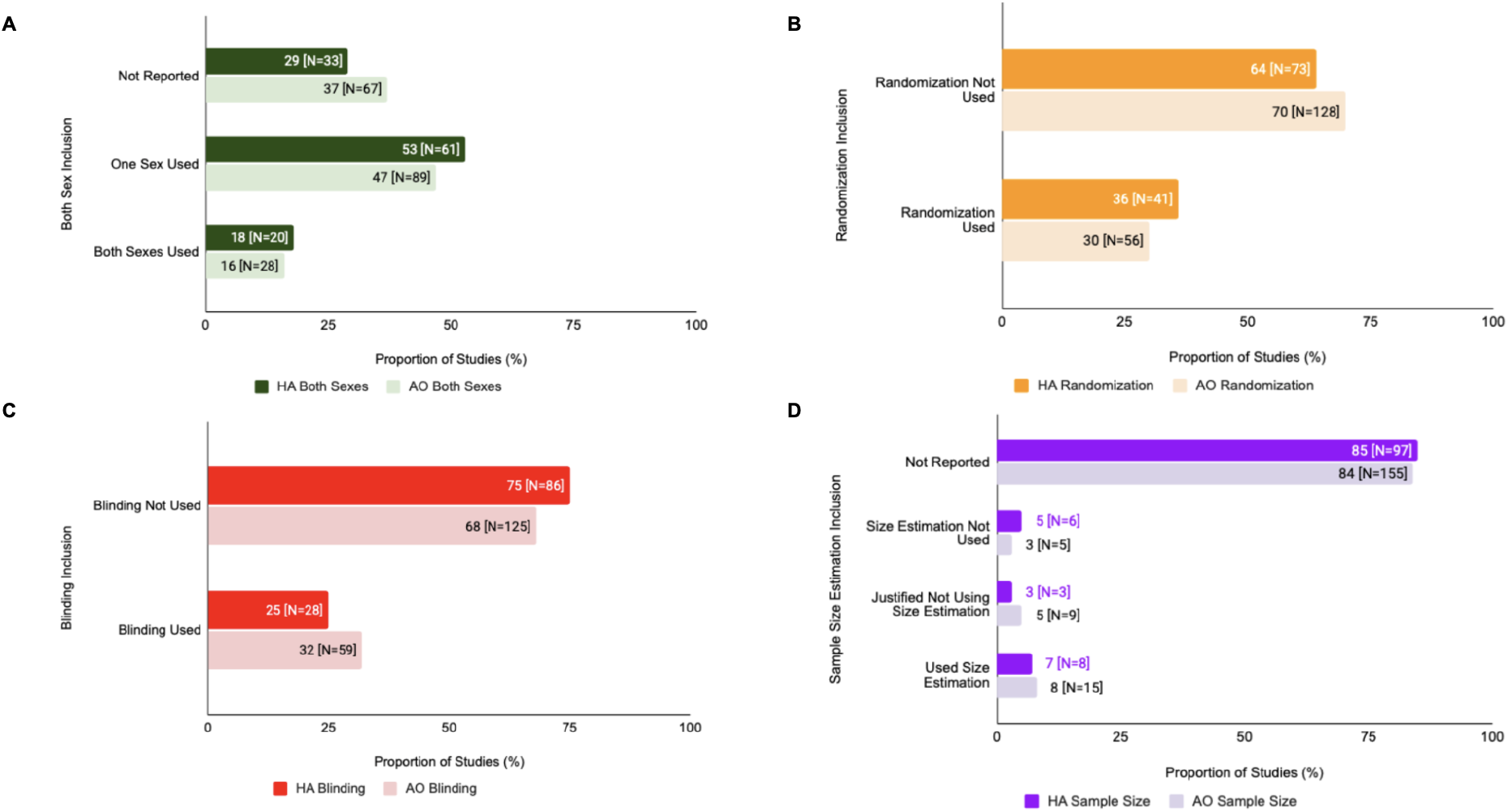
SDE inclusion varied in Human/Animal (HA) vs Animal Only (AO) studies. **(A) Both Sexes Analysis:** This figure depicts the inclusion of both sexes vs single sex in human/animal studies vs animal only studies (Chi-Squared analysis). **(B) Randomization Analysis:** This figure depicts the inclusion of randomization in human/animal studies vs animal only studies (Chi-Squared analysis). **(C) Blinding Analysis:** This figure depicts the inclusion of blinding in human/animal studies vs animal only studies (Chi-Squared analysis). (**D) Sample Size Analysis:** This figure depicts the inclusion of sample size estimation in human/animal studies vs animal only studies (Chi-Squared analysis).

However, there were no statistically significant differences in SDE reporting in human/animal or animal-only studies in terms of the sex of study subjects used (X^2^=1.775, df=2, corrected p=2.470), randomization (X^2^=0.7448, df=1, corrected p=3.104), blinding (X^2^=1.5715, df=1, corrected p=1.89), or sample size estimation (X^2^=2.2518, df=3, corrected p=2.609).

### Differences in SDE Reporting For Animals vs Humans within the Same Study

Within human/animal studies, there were variations in SDE reporting for human vs animal substudies (Figure 5). For human substudies in human/animal studies, 36% (N=41) used subjects of both sexes, 8.7% (N=10) used only one sex of study subjects, and 55.3% (N= 63) did not report the sex of study subjects used, as opposed to 17.5% (N=20), 53.5% (N=61), and 29% (N=33), respectively, for animal only studies. Randomization was used in 6% (N=7) of human substudies and 36% (N=41) of animal substudies, and blinding was used in 11% (N=13) of human substudies and 25% (N=28) of animal substudies in human/animal studies. In terms of sample size estimations for human substudies, 2% (N=2) performed sample size estimations, 12% (N=14) justified not using sample size estimations, 3% (N=3) did not perform sample size estimation, and 83% (N=95) did not report any information on sample size estimation. For animal substudies, these values were 7% (N=8), 3% (N=3), 5% (N=6), and 85% (N=97), respectively.

**Figure 5:**
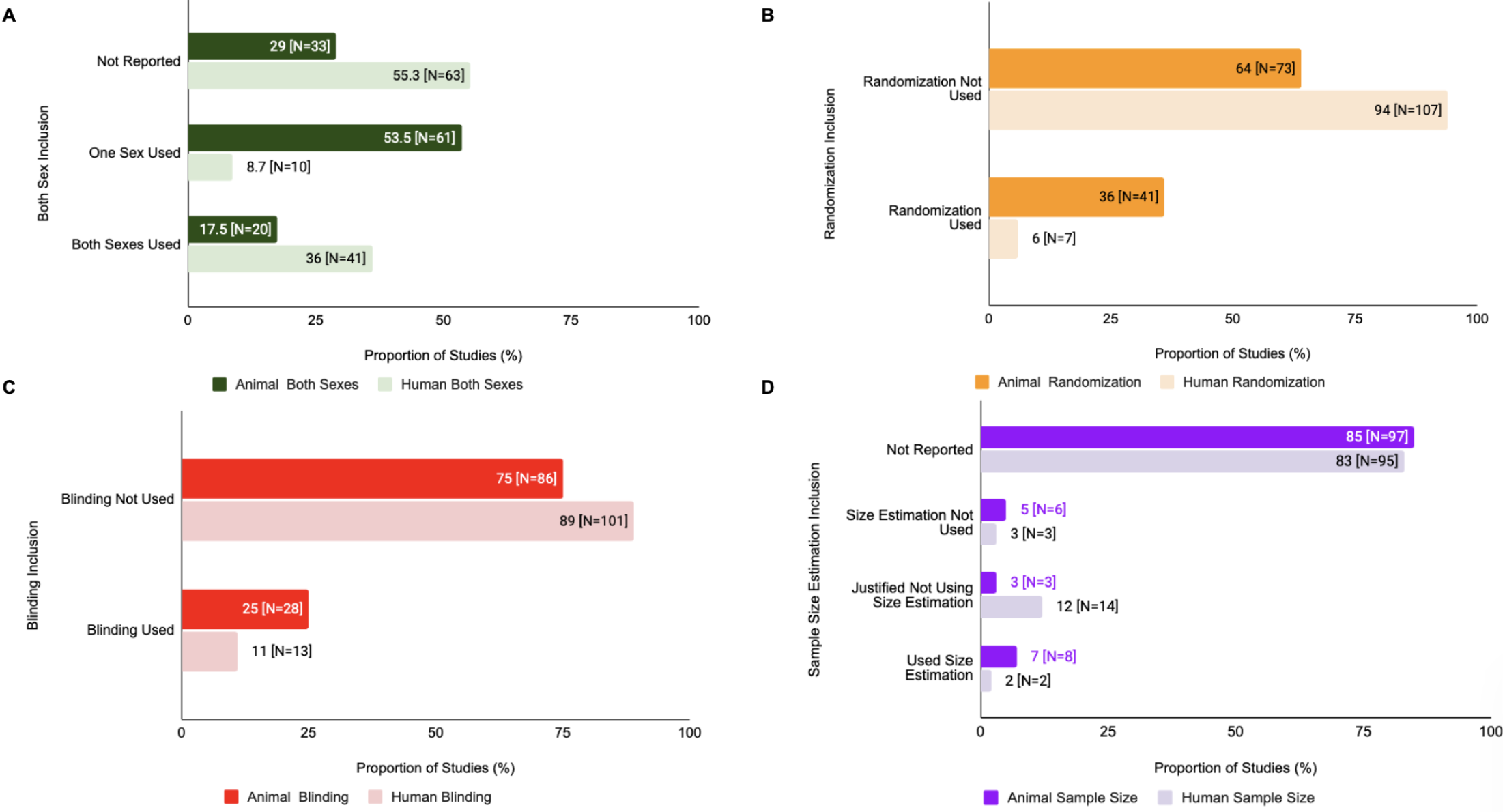
SDE inclusion for animal vs human substudies also varied within human/animal studies. **(A) Human/Animal Studies: Both Sexes Analysis**. This figure depicts the inclusion of both sexes vs single sex in human vs animal substudies in human/animal studies (Chi-Squared analysis). **(B) Human/Animal Studies: Randomization Analysis**. This figure depicts the inclusion of randomization in human vs animal substudies in human/animal studies(Chi-Squared analysis). **(C) Human/Animal Studies: Blinding Analysis**. This figure depicts the inclusion of blinding in human vs animal substudies in human/animal studies (Chi-Squared analysis). **(D) Human/Animal Studies: Sample Size Analysis** This figure depicts the inclusion of sample size estimation in human vs animal substudies in human/animal studies (Chi-Squared analysis).

There was a statistically significant difference in the use of randomization (X^2^=24.083, df=1, corrected p=1.85e-05) and sample size estimation inclusion (X^2^=38.911, df=3, corrected p=3.81e-07) of animal vs human substudies in human/animal studies. However, there was no statistically significant difference for blinding (X^2^=5.4878, df=1, corrected p=0.326) and sex of study subjects used in animal vs human substudies in human/animal studies (X^2^=3.123, df=2, corrected p=2.098).

### Differences in SDE Reporting in Different Animal Models

Reporting of biological sex SDE is greater in large animal studies whether including both sexes or stating only one sex was used when compared to small animal studies (Figure 6). Note that the number of small animal studies was far greater than the number of large animal studies that were used for this analysis. For small animals, both sexes were used in 16% (N=44) studies, a single-sex was used in 50% (N=135) studies, and sex was not reported for 34% (N=93) studies. Large animal studies exhibited a similar trend in that both sexes were used in 15% (N=4) studies, a single-sex was used in 58% (N=15), and sex was not reported in 26% (N=7) studies. There was no significant difference in the proportions of studies reporting sex between the small and large animals (chi-square test of interdependence, X^2^=0.689, df=2, corrected p=2.836). In small animal studies, randomization was used in 30% (N=82) whereas for large animal studies, 58% (N=15) studies used randomization. There was no significant difference in the proportions of studies using randomization between the large and small animals (chi-square test of interdependence, X^2^=6.995, df=1, corrected p=0.152). Small animal studies had 27% (N=74) of studies that used blinding whereas large animals had 50% (N=13) of studies that used blinding. There was no significant difference in the proportion of studies using blinding between the two variables (chi-square test of interdependence, X^2^=4.913, df=1, corrected p=0.390). Lastly, 86% (N=234) of small animal studies did not report any information regarding sample size estimation. For large animal studies, 69% (N=18) did not report any information regarding sample size estimation. There was no significant difference in the proportion of studies using sample size estimation between the two variables (chi-square test of interdependence, X^2^= 7.7154, df=3, corrected p=0.676). This data is summarized in Figure 6.

**Figure 6:**
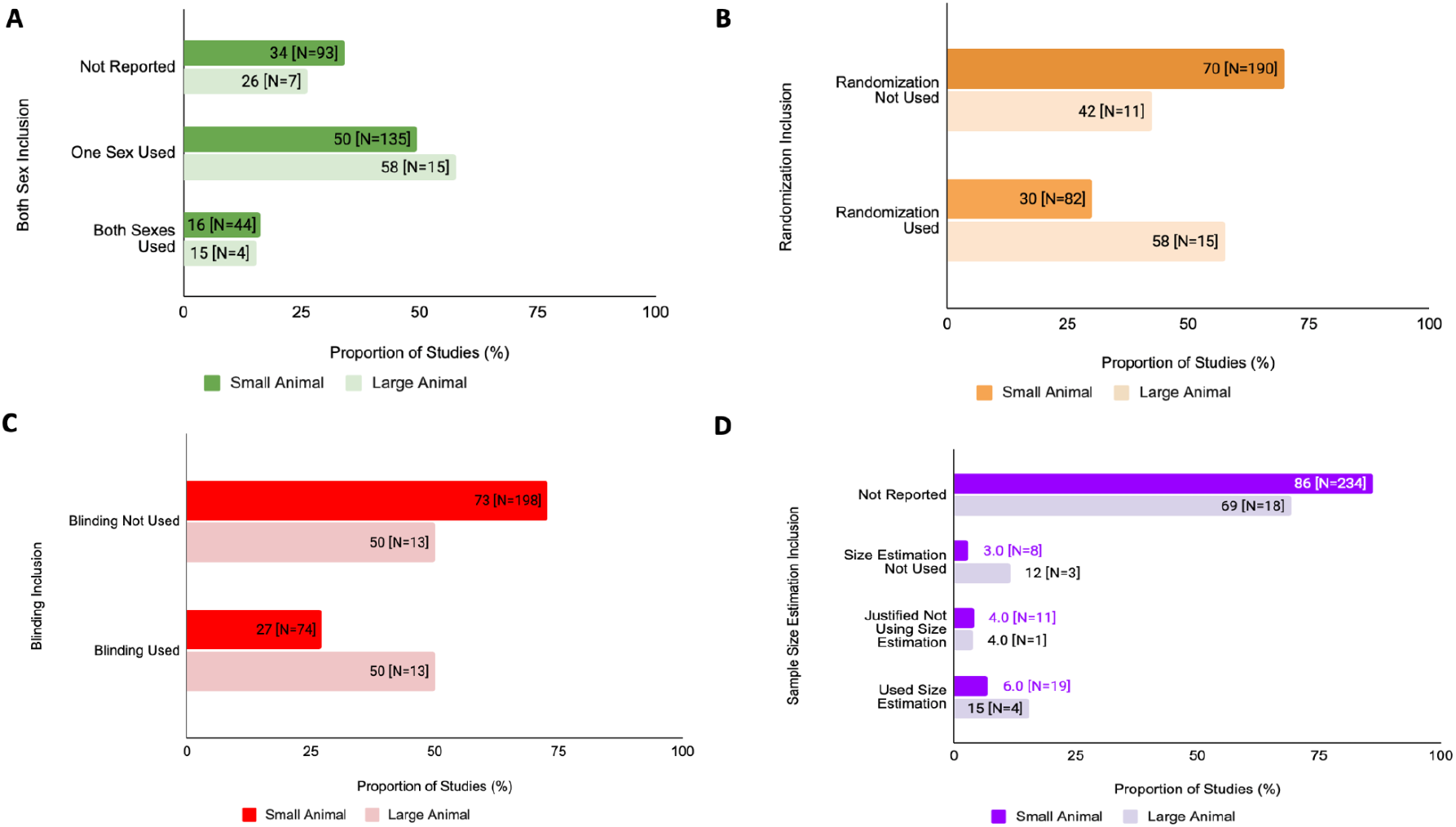
Proportion of SDE inclusion in preclinical cardiovascular studies differs between studies consisting of either small or large animal substudies from 2011-2021. **(A) Sex as a Biological Variable Analysis:** This figure compares the proportion of articles that reported incorporation of both sexes, a single sex, or omission of reporting for small or large animals (Chi-Squared analysis). **(B) Randomization Analysis:** This figure compares the proportion of articles that reported randomization for small or large animals (Chi-Squared analysis). (**C) Blinding Analysis:** This figure compares the proportion of articles that reported randomization for small or large animals (Chi-Squared analysis). **(D) Sample Size Estimation Analysis:** This figure compares the proportion of studies that reported use of sample size estimation for small or large animals (Chi-Squared analysis).

The proportion of single-sex studies was found to be greater for other animal models in comparison to rodents (Figure 7). Of the studies in which rodents were the primary animal model, 48.1% (N=128) were single-sex studies. This is less than other animal model studies where 68.8% (N=22) were single-sex studies. There is a statistically significant difference between single-sex inclusion in rodent studies and other animal studies (X^2^=4.073, df=1, corrected p=0.610).

**Figure 7:**
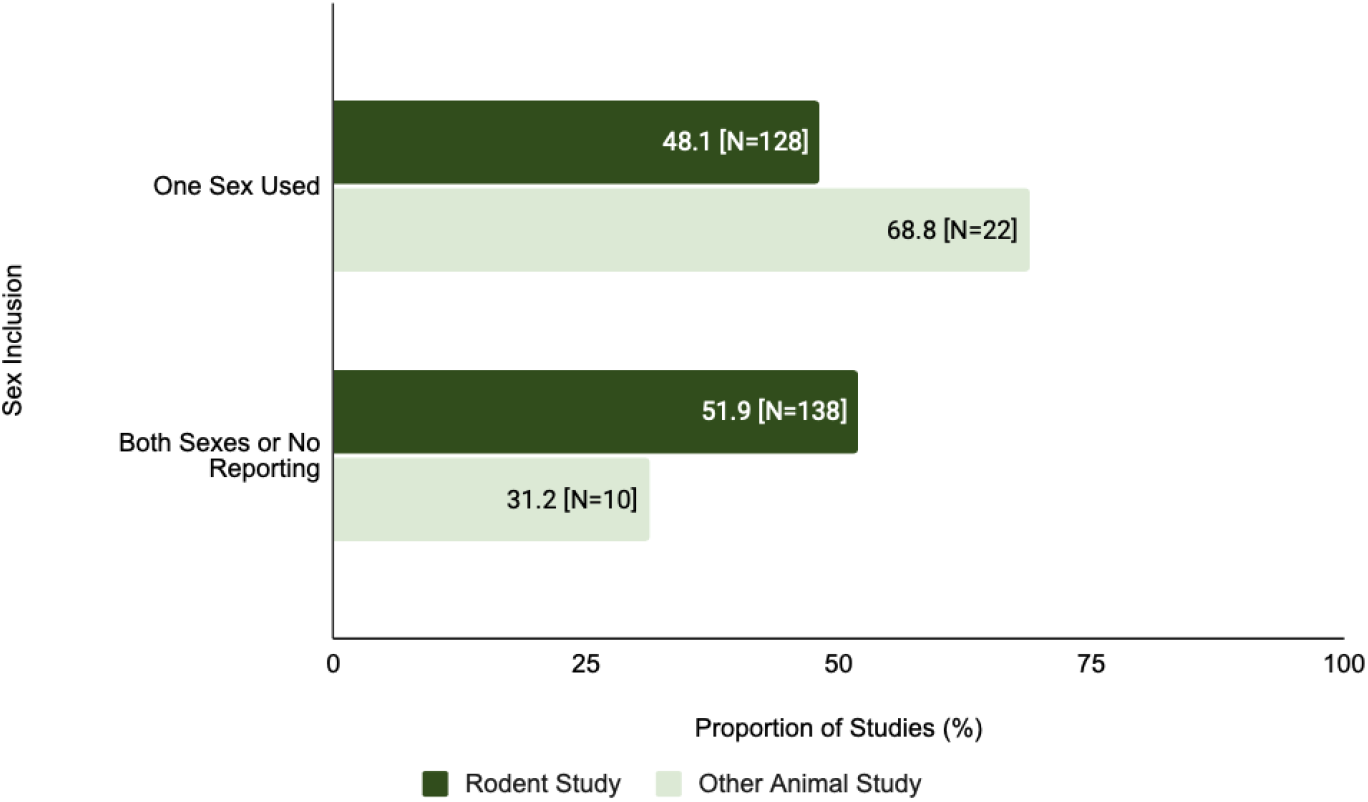
Sex inclusion in the study population varied in rodent vs other animal studies. This figure compares the proportion of rodent studies and non-rodent animal studies that were single-sex studies (Chi-Squared analysis).

## Discussion

We replicated and expanded a 2017 study by Ramirez et al. by assessing the reporting of four study design elements - inclusion of both sexes, randomization, blinding, and sample size analysis - in preclinical cardiovascular studies with either animal-only studies or studies with both human and animal substudies, over 10 years from 2011 to 2021. Overall, the inclusion of SDEs was low. 15.2% of animal-only studies included both sexes as a biological variable, 30.4% included randomization, 32.1% included blinding, and 8.2% included sample size estimation. Also, the incorporation of SDE in preclinical studies did not significantly increase over the ten years in the articles we assessed. Among human and animal substudies of human/animal studies, a significantly larger proportion of animal substudies reported usage of randomization and sample size estimation. Our conclusions serve as an informative checkpoint for the impacts of implementing ARRIVE (Animal Research: Reporting in In Vivo Experiments) and other protocols for enforcing methodological rigor (Lapchak et al., 2013; Ramirez et al., 2020; Williams et al., 2022).

### The Prevalence and Importance of Study Design Elements in Preclinical Research

The low reproducibility of preclinical studies may partially be attributed to the pressure to publish in high-impact journals, which is more easily achieved if a study has positive results (Mlinaric et al., 2017). Researchers are also incentivized to share positive results to secure funding or highly competitive academic jobs. Studies with limited SDE incorporation also make it harder to perform systematic reviews (O’Connor, 2018). Though the reproducibility crisis has been acknowledged by the NIH (Collins & Tabak, 2014) and mitigative efforts have been employed, the progress and success of these efforts continue to warrant assessment. In addition to Ramirez et al. (2017) finding that preclinical cardiovascular research studies had low SDE inclusion, Williams et al. (2022) determined that preclinical research articles do not adequately adhere to many ARRIVE guidelines, including having sufficient power for t-tests, use of randomization, and blinding. Our results support the findings of both of these previous studies.

### Study Design Elements

Although some SDEs can be relevant only to specific domains of a study, (Provencher, 2018), several SDEs can broadly be applied across preclinical research. The use of single-sex in a study compromises generalizability, therefore, the omission of sex as a variable may hinder later applications or the reproducibility of results in translational or clinical settings (Ramirez et al., 2017). Animal studies show a significant lack of incorporation of both sexes. In addition, there is a significant difference between the reporting of both sexes among rodent and non-rodent studies indicating that the type of animal model may be a factor in reporting (Figure 7).

Failure to apply randomization can exaggerate results, leading to unreliable findings (Hirst et al., 2014). This SDE is critical to preventing selection bias and is also the assumption upon which many statistical tests are based. In our study, the proportion of studies reporting the usage of randomization was low for both large and small animals.

The use of blinding in research also advantageously limits selection and procedural bias that could influence experimental results. Articles were screened for any reporting of blinding used whether it was single, double, or triple blinding, however, no significant differences were found.

Sample size estimation enables researchers to identify the necessary number of subjects for conclusions to be drawn and applied to the general population. This consists of predetermining a set number of animals needed for random group assignment. Sample size estimation was the lowest reported SDE among the four SDEs observed - only 8% of animal-only studies provided any justification, statistical or otherwise, for their sample size.

### Study Design Element Inclusion Over Time in the Past Decade

Although we hypothesized that the level of SDE inclusion would increase over time, the general prevalence of SDEs, whether they be applied to animal or human substudies, was remarkably low across the 298 articles screened (Figure 2). This contrasts with Ramirez et al. (2017), who found a positive trend in reporting for the journals they observed. It is important to note that their study screened studies published between 2006 and 2016, and we evaluated those published between 2011 and 2021. Another positive trend in temporal SDE inclusion in elements such as randomization and blinding has more recently been reported, however, sample size estimation and inclusion of both sexes remained low (Jung et al., 2021). In contrast, our findings indicate decreasing SDE inclusion rates for blinding and sample size estimation.

### The Influence of Animal and Human Subject Models on Study Design Element Inclusion

We evaluated SDE usage in animal-only studies compared to human/animal studies and SDE usage in animal substudies vs. human substudies within the same overall study. SDE incorporation in animal only and animal with human substudies were comparable though overall SDE reporting was low for both study types. Sample size estimation remained the lowest reported SDE for both study types.

Evaluating differences between SDE usage for animal or human subjects within the same study showed differences in the inclusion of randomization and sample size estimation. For these studies, a larger proportion of studies exhibit greater reporting of randomization and sample size estimation in animal substudies compared to human substudies.

### Study Limitations

Although all articles were subjected to randomized double screenings, we cannot be completely confident that an article did not use an SDE. In some instances, certain SDEs cannot be incorporated due to the conditions of the study, however, studies where SDE inclusion was not possible were still considered as lacking the SDE. Also, although the SDEs evaluated in this study are critical to conducting rigorous experimentation, other SDEs discipline-specific may ultimately be more relevant to the reproducibility of a paper.

### Future Directions

Future studies should expand the scope of SDEs reviewed in preclinical cardiovascular publications to include more domain-specific SDEs such as the use of comparison groups, units of concern, or other forms of subject allocation (O’Connor, 2018). A more detailed investigation of reasons publications do not include sample size estimation would also be extremely valuable, given the complexity of justification for including or not including the SDE. A further extension of this work would be to survey the authors of these studies to determine the underlying reason for either the omission or inclusion of different SDEs. This would allow both verification of this study’s screening, as well as expand upon the reasons why investigators do or do not use SDEs in their studies.

### Conclusion

The inclusion of SDEs improves reproducibility, which is important for translating preclinical findings into clinical outcomes. Contrary to our hypothesis, we found that over the past decade, there has been no significant increase in SDE incorporation in the preclinical cardiovascular publications we studied. Sample size estimation remains the least reported study design element. These trends all indicate the need for further efforts to increase the incorporation of rigorous study design elements in research projects to the point that they become routine. Future research efforts should evaluate a wider range of SDEs and investigate the reasons why SDEs have such low incorporation in preclinical cardiovascular research.

## Acknowledgments

The authors acknowledge support from the Stanford Program on Research Rigor & Reproducibility (SPORR). GCM acknowledges support from Kelsey Grinde, PhD from Macalester College for her guidance on the statistical analysis for this project.

